# Mind the gap: preventing circularity in missense variant prediction

**DOI:** 10.1101/2020.05.06.080424

**Authors:** Stephan Heijl, Bas Vroling, Tom van den Bergh, Henk-Jan Joosten

## Abstract

Despite advances in the field of missense variant effect prediction, the real clinical utility of current computational approaches remains rather limited. There is a large difference in performance metrics reported by developers and those observed in the real world. Most currently available predictors suffer from one or more types of circularity in their training and evaluation strategies that lead to overestimation of predictive performance. We present a generic strategy that is independent of dataset properties and algorithms used, to deal with circularity in the training phase. This results in more robust predictors and evaluation scores that accurately reflect the real-world performance of predictive models. Additionally, we show that commonly used training methods can have an adverse impact on model performance and lead to gross overestimation of true predictive performance.

## Introduction

Novel high-throughput sequencing technologies are used extensively for disease mapping, personalized treatment, and pharmacogenomics^1^. These next-generation sequencing (NGS) technologies identify so many missense variants that the interpretation of their effects has become an enormous challenge^2,3^.

To improve effectiveness of variant interpretation, computational approaches that can reliably distinguish benign from pathogenic variants are critical. The availability of more accurate classification methods will enable more effective integration of this genetic information into diagnosis, treatment and genetic counseling.

Many types of tools have been developed that predict the pathogenicity of missense variants. Some use features describing different aspects of evolution, biochemical properties of amino acids and protein structure^4,5^ In an attempt to improve on those predictors, so-called meta-predictors were developed^6–17^, where these predictors use the predictions of individual predictors as features. However, the real clinical utility of current approaches is rather limited, and a large difference remains in performance metrics reported by developers and those observed in the real world^18–22^

Comparing real-world performance of pathogenicity predictors is a daunting task. As described by Grimm et al.^23^, the evaluation of predictors is hindered by different forms of circularity. Circularity arises when (1) the same variants are used in the training and evaluation set, or (2) variants from the same genes are used in the training and evaluation set, both leading to overestimation of real-world predictive performance.

Where Grimm et al. mainly focused on assessing the impact of different types of circularity on the comparative evaluation of predictive tools, we show that these concepts also apply for the development of novel predictors. Even more, we show that failing to deal with circularity in the development phase is a major cause for the observed gap between reported performance metrics and those subsequently observed in real world scenarios.

Developing a predictive tool usually involves several iterations of data selection, feature selection, model parameter tuning and cross-validation experiments guided by validation scores as a metric of predictive power.

This process is subject to circularity if training, evaluation and test sets are not properly constructed. It is well-accepted that using the same variants in training and validation sets (type 1 circularity) leads to overfitting and overestimated performance. However, the drawbacks of including variants from the same genes in both the training and validation sets are less well known and not immediately obvious. With some exceptions^5,24^, current state of the art predictors^6,25^ are trained resulting in embedded circularity. We show that this type of training regime leads to overly optimistic performance metrics.

We present a training strategy that eliminates both type 1 and type 2 embedded circularity. The main feature of this strategy is the rigorous separation of genes between training, evaluation and test sets. By applying an extensive “k*1 fold” training architecture, we show that ignoring circularity effects in the training phase might actually result in less predictive power. In all cases not adjusting for type 1 and type 2 circularity leads to grossly overestimated real-world performance. We demonstrate that our strategy produces classifiers with the best real-world performance, independent of algorithms used. In addition, this strategy results in evaluation scores that are representative of real-world predictive performance.

## Methods

### Definitions

#### Sample

A single observation or record consisting of a number of features that describe the observation.

#### Dataset

A collection of data consisting of individual samples, with a label associated with each of the samples.

#### Training set

A dataset that is used for training a machine learning algorithm. The algorithm will generally observe every one of the samples in the dataset directly and attempt to find a relationship between the features and the labels.

#### Validation set

A dataset that is not observed directly by a machine learning algorithm simultaneously with a given training set. This dataset is used to validate the performance of the machine learning algorithm by observing prediction performance during training without attempting to learn a relationship between the features and the labels. The validation set may be used to stop training at some point in order to avoid overfitting.

#### Test set

A dataset that is used to assess the performance of a final model in machine learning. This dataset is not used to regulate training in any way and only serves to evaluate the model after training.

### Datasets and data preprocessing

The reference variant dataset used in this manuscript was composed of variants from ClinVar^26^ and gnomAD^27^. ClinVar variants were included if the Clinvar review status was one star or higher, excluding variants with conflicting interpretations. ClinVar variants with “Benign” and “Likely benign” interpretations were included and labeled as benign variants, whereas those variants with “Pathogenic” and “Likely pathogenic” interpretations were included and labeled as pathogenic variants. gnomAD variants were selected from the set of genes obtained from ClinVar, and labeled benign if the minor allele frequency exceeded 0.1%. This resulted in a dataset with 147497 distinct variants, 27.7% of which were labelled pathogenic.

Features describing the individual variants were obtained from 3DM^28^. Features include alignment-derived metrics such as amino acid conservation and sequence entropy, as well as overall alignment descriptors such as the number of species observed and alignment depth. Where protein structures were available, structure-derived properties were included, such as surface-accessible areas, secondary structure information, and the number of hydrogens bonds a particular residue is involved with.

To assess the impact of training regimes on the performance of meta-predictors a separate experiment was performed. A distinct dataset was compiled, where predictors scores are used as features describing the reference variants Predictor scores that were used to train a meta predictor classifier were obtained from dbNSFP^29,30^. The predictors were chosen to mimic the constituent predictors of REVEL. Predictors used were FATHMM^31^, PROVEAN^32^, Polyphen2^5^, SIFT^33^, MutationAssesor^32^, MutationTaster^34^, LRT^35^ and GERP++^36^, in combination with statistics from several alignments, specifically: SiPhy^37^ 29-way log odds, phyloP20 mammalian odds, phyloP7 vertebrate odds, phyloP100 vertebrate odd, phastCons 20-way mammalian odds, phastCons 7-way vertebrate odds and phastCons 100-way vertebrate odds.

### Cross-validation dataset creation

*k*l* cross-validation^38^ was used to train and evaluate model performance. A nested strategy was applied to generate training, validation and test sets from the full dataset. First, the full dataset is stratified into *k* partitions. In *k* iterations each partition is used as an outer test set.

Ten outer test sets were created with the group *k*-fold principle: a dataset is split into *k* partitions, each of which contains a unique set of samples belonging to genes that do not occur in any of the other partitions (see stratification below). In *k* iterations, *k*-1 partitions are selected as a set that was later used to construct the training and validation set, while the leftover partition is used as a test set (figure 1).

**Figure 1.**
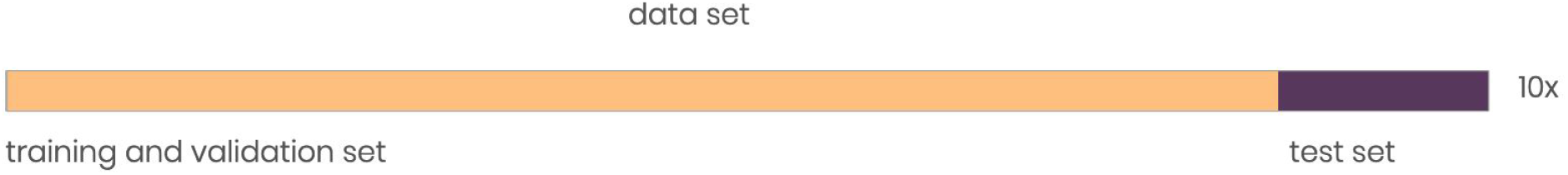
The 10 test sets used in this study are created by dividing the data set into 10 partitions, where 9 partitions are used for training and validation (inner set), and 1 partition is used for testing model performance (outer set). The test sets exclusively contain samples belonging to genes that do not occur in the corresponding training and evaluation sets.

The remaining *k* – 1 partitions are then combined to form the inner set. In each of the *k* iterations, the inner training set is divided into *l* partitions. In *l* iterations each of the partitions is picked once as a validation set, while the remainder is combined to form the training set (figure 2). This results in a total of *k*l* folds, each with a training, validation and a test set.

**Figure 2.**
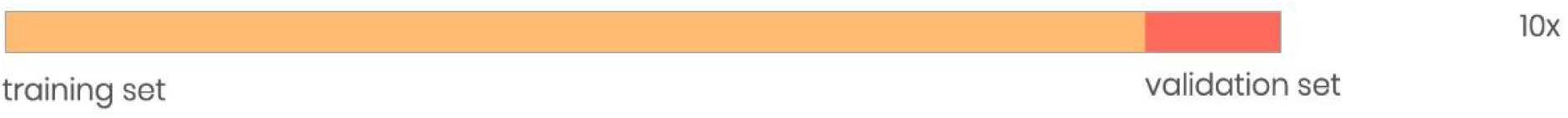
The inner set is used to create 10 pairs of training and validation sets.

Within each fold, a variant can be present only once, in either the training, validation, or test set. To assess the effect of the training and evaluation strategies, we constructed training and evaluation sets in three different ways, where the outer test sets were identical in all scenarios described.

#### Gene split

training and validation sets are constructed in such a way that validation sets exclusively contain samples belonging to genes that do not occur in the corresponding training set.

#### Position split

training and validation sets are constructed in such a way that validation sets exclusively contain samples that do not occur at the same protein position. The training sets contain samples belonging to genes that also occur in the validation sets. E.g. for a given position in a gene there might be multiple variants described. In this position split scenario all the variants on that position end up either in the training set or the validation set.

#### Random split

training and validation sets are constructed by distributing variants randomly, irrespective of position or gene.

Applying the approach described above resulted in three datasets for three experiments:

1. Outer split: gene (10 folds), inner split: gene (10 folds), total 100 folds
2. Outer split: gene (10 folds), inner split: position (10 folds), total 100 folds
3. Outer split: gene (10 folds), inner split: random (10 folds), total 100 folds

The impact of different data splitting strategies on unknown genes in a cross validation scenario can be discerned, since the 10 outer folds are identical across experiments.

Model performance on the test set containing unseen genes is used as an indicator of real world predictive performance. When model performance is evaluated on genes that the model has seen, the class distribution in the training set is far more likely to reflect the distribution of the test set than the true probability of deleteriousness (see *Stratification).* This means that the test set is biased towards good performance for the model, instead of reflection of the real world. When testing on unseen genes no such bias is present, which makes it more suitable for the evaluation of model performance in a real world scenario.

### Stratification

Stratification describes a process whereby each partition in the *k*-fold strategy receives an approximately equal number samples belonging to a specific group. The groups in this case are the class imbalance bins that each gene belongs to. By distributing the genes as fairly as possible according to these bins across all the partitions, a partitioning is achieved that approaches the lowest possible inequality between folds while preserving the gene-grouping constraint.

Class imbalances between training and validation sets can result in misleading performance statistics. Ideally, every partition has a similar class balance, as this results in training and validation sets with a similar class balance. We therefore place extra constraints on the split to reduce inequality between the mean classes in each partition.

For each gene we find the class balance, expressed as a real number between 0 and 1. This class imbalance is then multiplied by a factor N and round to an integer. This effectively bins the class imbalances into different groups. We use this binned class imbalance to add a stratification constraint to the *k*-fold partitioning logic.

### Normalization

Neural networks train faster and perform better when data is normalized to have a mean of 0 and a standard deviation of 1^39^. To avoid errors introduced by normalizing the training set and the testing set separately, all feature data was normalized based on data generated for the entire proteome. By generating means and standard deviations for the entire dataset upon which inference would take place we ensure that normalization is representative for future samples and not biased by the distribution of labeled records. Tree-based learning methods are not affected by data normalization.

### Model training

Model training was done using the Python versions of the models described. Several different model types were used to cover multiple scenarios where using different training regimes can affect performance.

### NODE

Neural Oblivious Decision Ensembles^39^ represent the state of the art in Neural Network learning models specialized in classifying tabular data. In their paper Popov et al. demonstrate that NODE outperforms gradient descent boosted trees (GDBT) in all scenarios with default hyperparameters.

This makes it an attractive candidate for inclusion. NODE was used with default hyperparameter settings.

### Random Forest

The Random Forest^40^ learning algorithm uses an ensemble of decision trees in order to be more robust against overfitting than single decision trees. It is an algorithm used often in scientific applications.

We use the Random Forest technique in two instances. First with our own features and a sensible set of hyperparameters: 1000 estimators, gini coefficient splitting and the maximum fraction of features set to 1. Second with a reproduction of REVEL’s features and hyperparameters to demonstrate how different splits impact the metaclassifier paradigm.

### CatBoost

CatBoost^41^ was picked as a representative Gradient Boosting method, as it constitutes the state of the art in Gradient Boosting. Only one other predictor in use today (ClinPred^16^ uses XGBoost) uses this method, but the ubiquity of Gradient Boosting in other prediction problems and its outstanding performance makes this a logical next step in the evolution of missense variant prediction. We show that the problems we encounter also pertain to this method of predictions.

Gradient boosting methods are extremely effective learners^42^, able to fit a dataset to a great extent. The number of iterations is set to 50000, with a learning rate of 0.1 in order to ensure that the algorithm converged. The metric to be optimized was set to LogLoss. Finally, GPU training was used to speed up training, which resulted in the max_bin parameter to be set to 254.

### Optuna

Hyperparameter tuning was used to build Random Forest models to simulate what happens when the validation set is used to optimize hyperparameters. The Optuna library^43^ was selected to perform optimization using its standard bayesian approach^44^. We used Optuna to run automated hyperparameter optimization with the Random Forest algorithm using a subset of the k*l fold dataset: 5 outer folds and 5 inner folds in k*l cross validation. This subset was used to decrease optimization time. Optimal parameters for each of the training regimes were selected based on the validation score.

### Matthew’s Correlation Coefficient

Matthew’s Correlation Coefficient (MCC)^45^ was used to evaluate model performance. It is a measure used in (binary) classification that gives a good sense of classifier performance, even when the evaluation dataset is imbalanced^46^. The measure returns a value from −1 to 1, with 1 indicating a perfect score, 0 indicating prediction performance that is no better than random and −1 indicating the inverse of a perfect score, with every classification being the opposite of the true label. This results in interpretable scores independent of the class balance.

## Results

We evaluated the performance of the trained models on the test sets, which contain variants in genes to which these models were never exposed. We used three different types of machine learning algorithms (NODE, CatBoost, RandomForest) that were trained on the dataset with in-house features. We also used the methodology of REVEL to assess the effects of different training regimes for metaclassifiers, both in a static scenario as well as with a hyperparameter search using bayesian optimization. Each split method (gene, position, random) was tested 100 times, resulting in 300 measurements.

Important to note is that a small difference between validation performance and test performance is ideal; in that case the expected model performance is a clear indicator of real world performance, and no circularity is introduced. If, however, there is a large gap between validation performance and test performance, the model is able to achieve excellent predictive performance on the validation set, but is not able to achieve the same level of performance in the real world. In that case, there is a clear overestimation of model performance due to introduced circularity.

The focus of this manuscript lies with assessing the impact of different training regimes on real world predictive performance in the context of different predictors. Therefore, in the figures below the effects of using different training and validation sets and the relative impact of the performance on the test set are visualised, instead of comparing absolute MCC scores across different scenarios.

When using NODE we see that the gene split validation and test results are very close to each other (figure 3). There are no statistically significant differences between the performance scores (p = 0.26, two sided Student’s T-Test). Here, the validation scores for the best model (based on gene split) accurately reflect the real world performance. In the case of the position and the random split we notice two phenomena: there is a large gap between the validation performance and the test performance, and the test performance is lower than the test performance for the best model that was trained using the gene split approach. Results for all prediction methods (including NODE) are summarized in figure 4.

**Figure 3:**
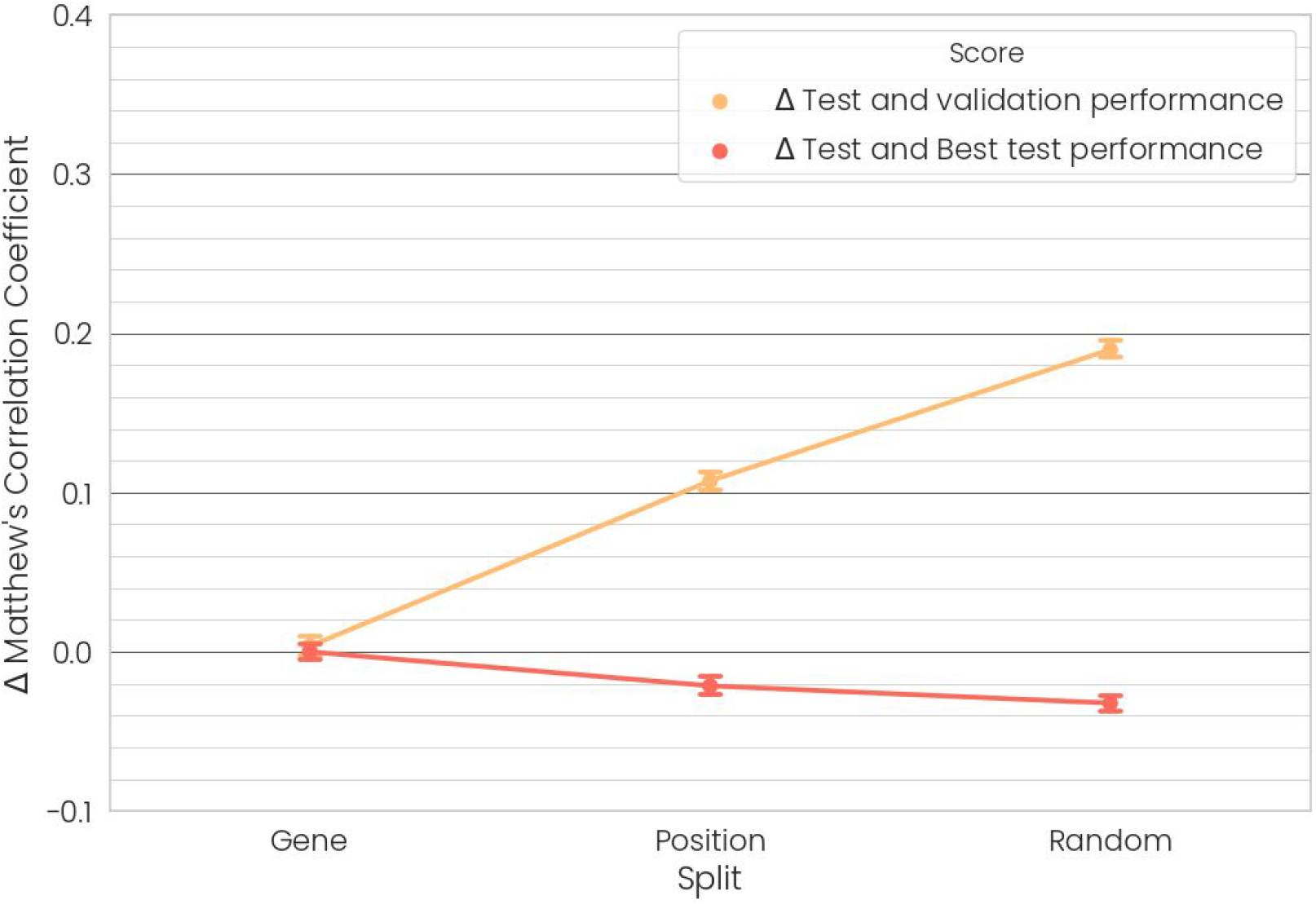
Performance changes in NODE when evaluating on 100 (10*10) k*l fold iterations. Performance on the test set decreases while performance on the validation set increases drastically.

**Figure 4:**
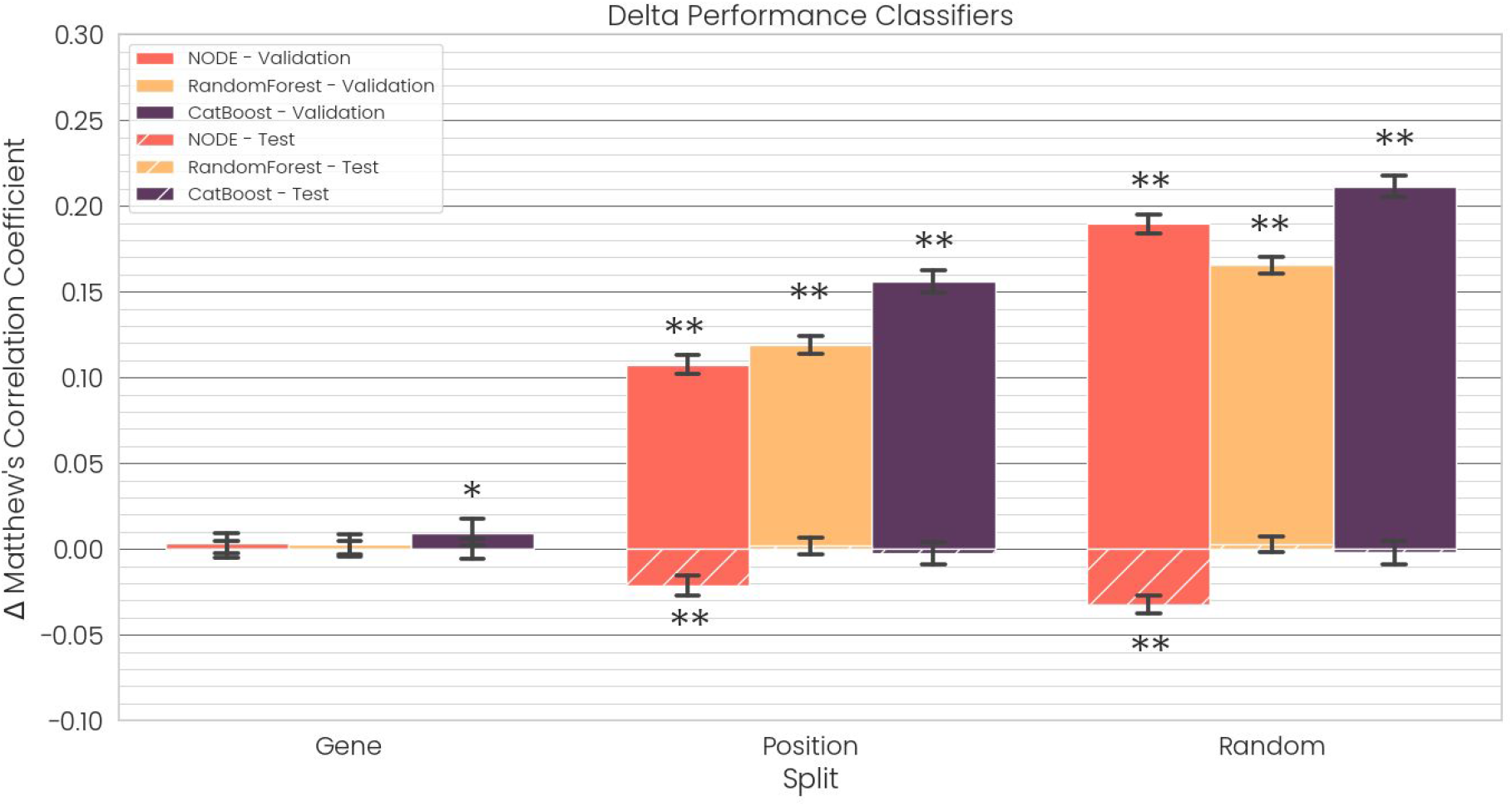
Summarized results for delta performance. Solid bars represent the difference in MCC between the test and validation set. Hatched bars show the difference in performance on the test set relative to gene split performance. Error bars denote the 95% confidence interval. One asterisk indicates a significant difference (p<0.05), two asterisks indicate very significant differences (p<0.005)

With CatBoost we observe very similar effects of using different split methods. In figure 4, we see that the gene split validation and test results are very close to each other. There is no statistically significant difference between the performance scores (p = 0.11, two sided Student’s T-Test). Here, the validation scores of the gene split based models accurately reflect the real world performance. There is a large gap between the validation performance and the test performance for both the position split and the random model.

Due to the ability of tree based methods to resist overfitting, the test scores do not deteriorate in the position and random regimes (no statistical difference from the gene split, p = 0.29, two sided Student’s T-Test). This means that the real world performance of any tree based model is likely to be very similar. However, when not using a gene separation approach, the validation score is not representative of real-world performance but is bound to be a great overestimation.

In order to investigate whether the introduced circularity effects were independent of the features used to describe variants, an effort was made to emulate the methodology of meta predictors like REVEL^6^. This involved training a Random Forest model on a set of predictions from other models as features. When working with a technique that does not use early stopping or validation set monitoring, the test set performance remains statistically the same as model parameters are not directly reliant on the validation set. However, we still find that the validation scores improve dramatically when different splits are used. The gene split scores are not significantly different between the validation and test sets (Student’s t-test, p>0.13). Validation scores reported based on position or random splits are thus inflated.

We applied automatic hyperparameter tuning to the Random Forest using Optuna. Pearson correlation analysis of the results obtained from 100 rounds of optimization on the random and position split show a negative relation between validation and test performance (−0.52 and −0.55 correlation respectively, p <<< 0.05). Optimizing the gene split resulted in a pearson correlation of 0.998 between test performance and validation performance (p <<< 0.05). These optimization results are illustrated in figure 5. This means that optimizing on a gene split will lead to a model that performs better on the independent test set, whereas optimizing on other splits is likely to lead to worse performance on the test set. Figure A.2 in the appendix shows the distribution of the test runs.

**Figure 5:**
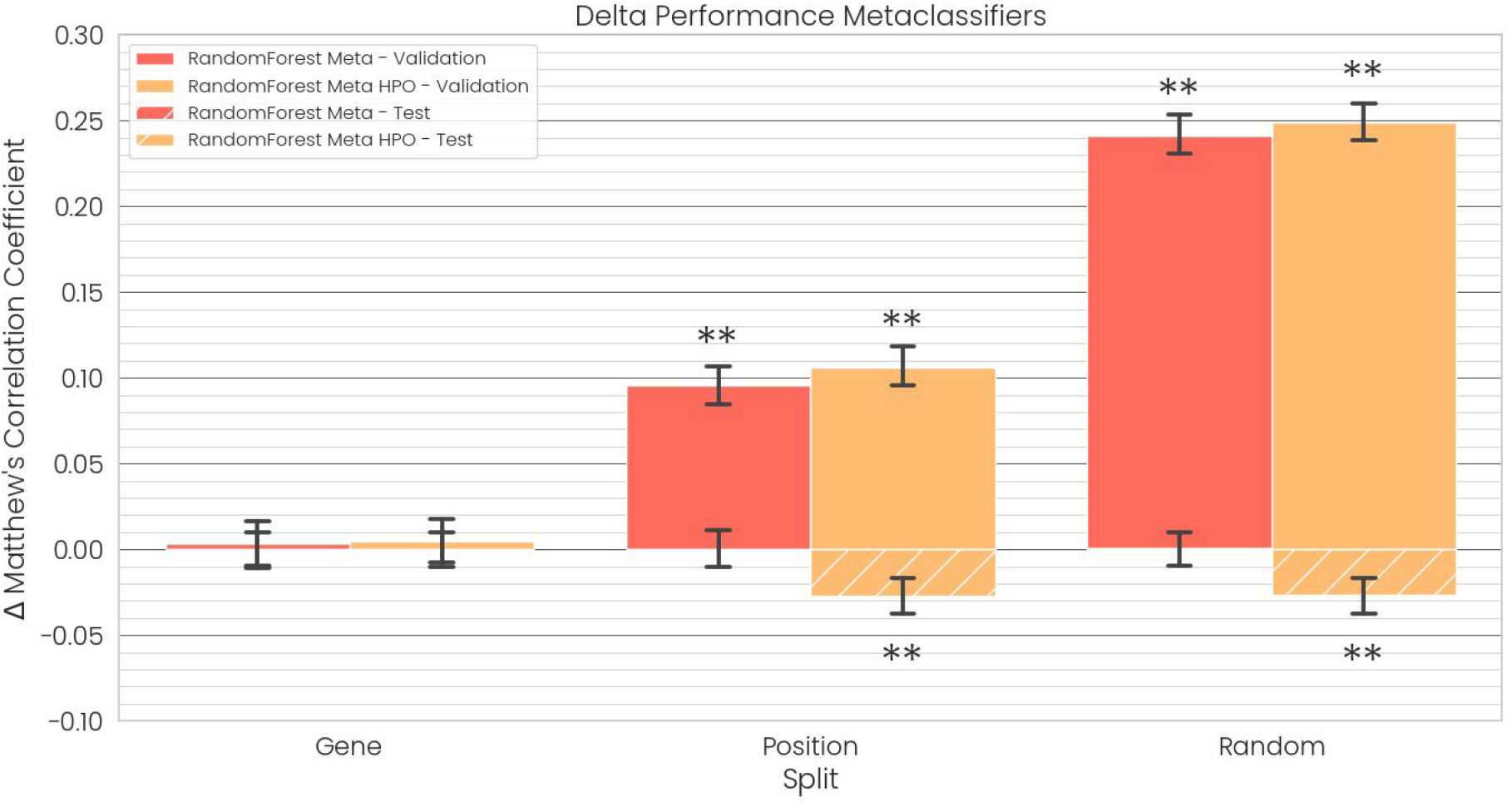
Summarized results for all the metaclassifiers. Solid bars represent the difference in MCC between the test and validation set. Hatched bars show the difference in performance on the test set relative to gene split performance. Error bars denote the 95% confidence interval. One asterisk indicates a significant difference (p<0.05), two asterisks indicate very significant differences (p<0.005)

When these parameters are run in the full k*l fold cross validation scenario, we find that models with the optimal hyperparameters for random and position splits yield significantly reduced performance compared to the gene split. It also increases the delta between validation performance and test performance. Hyperparameters selected by optimizing on the gene split resulted in better test performance and no significant difference between the test and validation performance (p=0.47). This is illustrated in Figure 5.

### Impact on training time

Depending on the number of samples and the learning algorithm chosen, training a model can take anywhere from minutes to days. Early stopping can stop training a model before it overfits. Random and position splits yield validation sets that closely reflect the training set, which results in more iterations needed before the validation loss stops dropping. When using the gene split learning often stops far earlier into the training process. Less iterations means less time spent training, with no drawbacks in terms of model performance. The training time improvement is illustrated in figure 6. Using a gene split results in a 4-7x speed up, with better or equivalent prediction performance.

**Figure 6:**
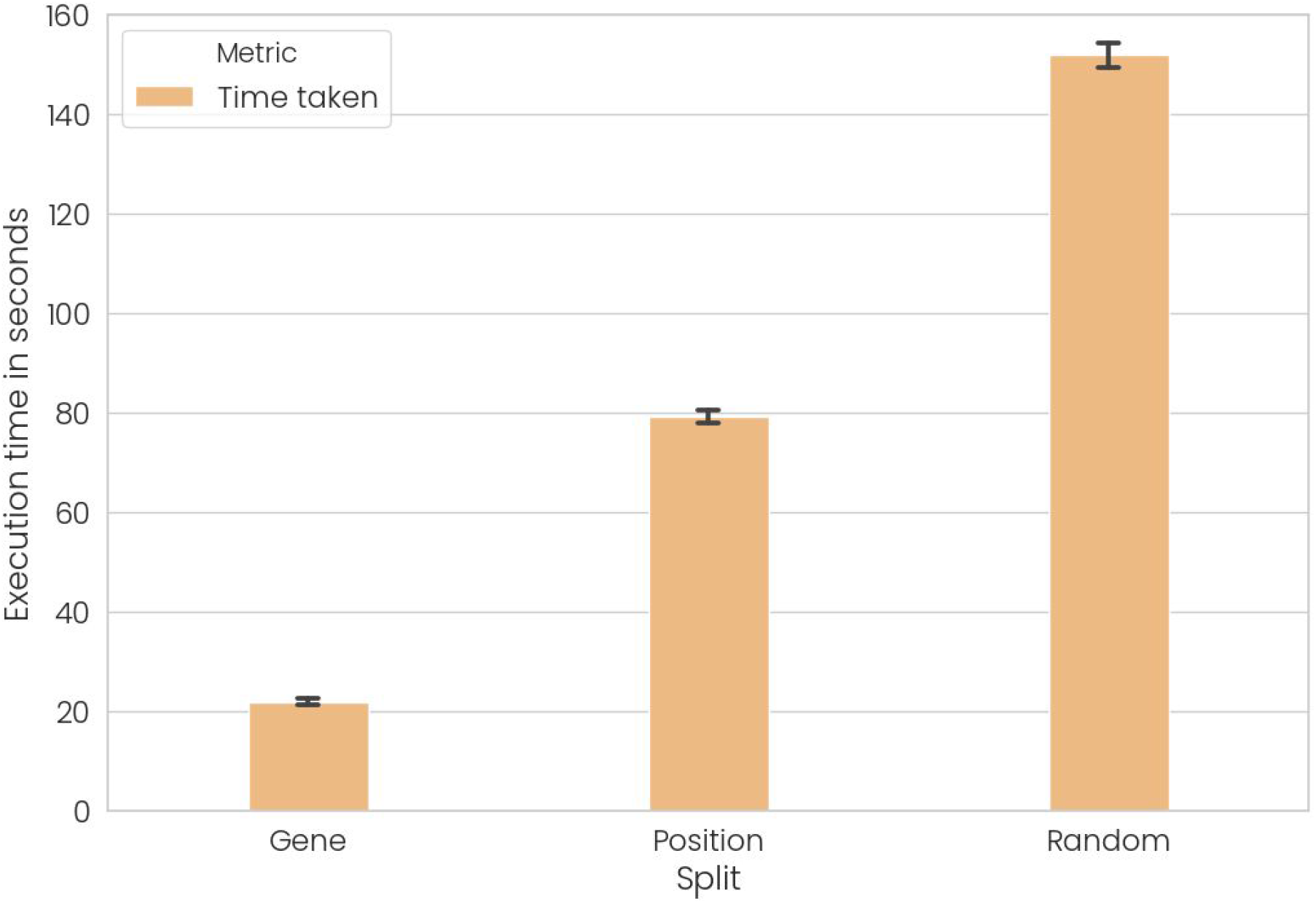
Average time in seconds needed to train one fold in CatBoost until convergence with different splits. Using a gene split results in a 4-7x speed up, with better or equivalent prediction performance.

## Discussion

We performed a comparative evaluation of the impact of different training regimes for pathogenicity prediction tools, and demonstrated that cross-validation scores are only indicative of real world performance when circularity is actively avoided. This is independent of training features and the machine learning algorithms used. We showed how and to what extent ignoring these effects could lead to overly optimistic assessments of tool performance.

The primary reason for the fact that the use of different validation datasets produce different test scores seems to be overfitting. Validation sets do not directly inform the training procedure, but depending on the machine learning method they do affect the overall training process. In Neural Networks and gradient boosting they determine the amount of iterations that are run on the dataset. With machine learning techniques that do not use a validation set directly they influence the model search by the researcher, which results in different model parameters. As a rule, training is continued until classifier performance on the validation set stops increasing. In practice, we see that position and random splits allow for much more training iterations before validation performance plateaus than a gene level split. Even with similar datasets, it seems that after a certain amount of training the classifier starts to extract phenomena that take advantage of the distribution inside genes, instead of learning to extract knowledge that can be extended to other, unseen genes.

### Recommendations

We recommend that novel predictors be evaluated by reviewers with these issues in mind. As we have shown in this work, reported cross-validation performance metrics are close to meaningless when training strategies are applied that do not properly take circularity into account. In the case of our metaclassifier replica, for example, our validation results suggested a score 0f 0.88 MCC, while the true test score was 0.64 MCC. We would like to suggest that novel prediction tools make their training datasets available on publication (including cross-validation splits), as both type 1 and type 2 circularity cannot be excluded if any portion of a training dataset is kept private. This would both benefit the evaluation of how well tools generalise to real-world scenarios, as well as enable fair comparisons between predictors. In general, it would mean a much needed increase in reproducibility and transparency of the field.

## Conclusion

Not properly taking circularity into account during training is harmful in three ways: First, it misleads the user into believing that the real world performance of their model is far better than it actually is. Second, it can result in an objectively worse model in practice. This is also true when (automatic) hyperparameter optimization occurs. Finally, when datasets are not separated by genes, training time is lengthened without yielding any performance increase. Instead, by using a gene split approach it is made sure that validation scores are close to scores that are expected on an independent test set created from unseen genes. Additionally, this results in the added benefit of decreased training times. We therefore recommend that missense variant predictors use only stringent gene split approaches to evaluate their performance as a matter of methodological rigor and as a guideline for improving predictor performance. These recommendations were taken into account in the development of Helix, a novel pathogenicity predictor (manuscript in preparation).

## Acknowledgements

We thank Prof. Dr. Gert Vriend for critically reviewing this manuscript.

(Part of) The research leading to these results has received funding from the European Union’s Horizon 2020 research and innovation programme under grant agreement No 634935 (BRIDGES), No 635595 (CarbaZymes) and No 685778 (VirusX).

## Appendix

## Results table

The results for each of the classifiers with different splits are reported in table A.1 with associated statistical significance.

**Table A.1:** Summarizes results for the different classifiers that were tested. One asterisk indicates a significant difference (p<0.05), two asterisks indicate very significant differences (p<0.005)

| Classifier | Split | $\Delta$ Validation/Test | $\Delta$ Test gene split |
| --- | --- | --- | --- |
| NODE | Gene | -0.003 |  |
|  | Position | -0.107 ** | -0.021 ** |
|  | Random | -0.189 ** | -0.032 ** |
| RandomForest | Gene | -0.003 |  |
|  | Position | -0.095 ** | -0.0002 |
|  | Random | -0.241 ** | -0.0005 |
| RandomForest<br>Hyperoptimization | Gene | -0.004 |  |
|  | Position | -0.106 ** | -0.026 ** |
|  | Random | -0.248 ** | -0.026 ** |
| CatBoost | Gene | -0.009 * |  |
|  | Position | -0.156 ** | -0.002 |
|  | Random | -0.211 ** | -0.001 |

### Hyperparameter optimization results

**Figure A.1:**
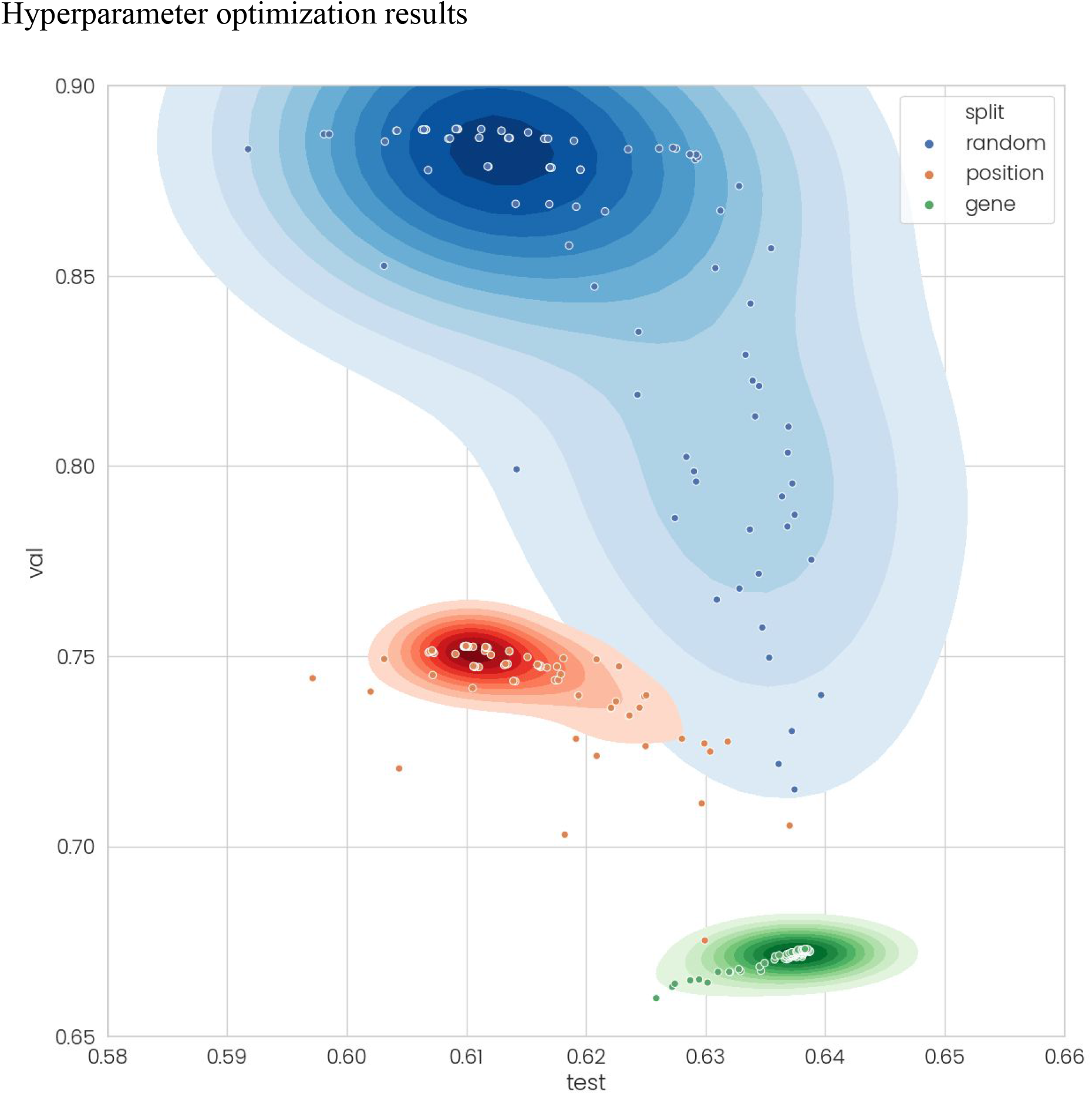
When hyperparameter optimization is applied to a robust algorithm like Random Forest, the choice of validation set is of utmost importance. In the random case, models which work best on the validation set yield poor performance on the test set. Hyperparameter optimization of the gene split set resulted in optimal models with better test scores. The shaded contours show the density of each of the distributions.

